# A Connectome-wide Functional Signature of Trait Anger

**DOI:** 10.1101/2020.10.14.338863

**Authors:** M. Justin Kim, Maxwell L. Elliott, Annchen R. Knodt, Ahmad R. Hariri

## Abstract

Past research on the brain correlates of trait anger has been limited by small sample sizes, a focus on relatively few regions-of-interest, and poor test-retest reliability of functional brain measures. To address these limitations, we conducted a data-driven analysis of variability in connectome-wide general functional connectivity, which has good test-retest reliability, in a sample of 1,048 young adult volunteers. Multi-dimensional matrix regression analysis showed that individual differences in self-reported trait anger maps onto variability in the whole-brain functional connectivity patterns of three brain regions that serve action-related functions: bilateral supplementary motor area (SMA) and the right lateral frontal pole. Follow-up seed-based analysis confirmed that high trait anger is associated with hyperconnectivity between these three regions and the somatomotor network as well as hyperconnectivity and hypoconnectivity between SMA and default mode and visual networks, respectively. Supplementary targeted analyses based on theoretical and empirical grounds further revealed that high trait anger is associated with hyperconnectivity between the amygdala and dorsomedial prefrontal cortex, dorsal anterior cingulate cortex, and striatum. These patterns suggest that the dispositional tendency to more easily experience frustration and anger is associated with variability in the functional connectivity of brain networks supporting somatomotor, affective, self-referential, and visual information processes. The emergence of action-related brain regions from our connectome-wide analysis is consistent with trait anger as reflecting a greater propensity to provoked action.

---

Trait anger refers to an individual’s dispositional tendency to more easily experience frustration, resulting in a decreased threshold for feeling angry in a wide range of situations (Spielberger, 1991). High trait anger has been broadly linked with both behavioral outcomes including higher reactive aggression and violent behavior (Bettencourt, Talley, Benjamin, & Valentine, 2006; Eckhardt, Jamison, & Watts, 2002) and health outcomes including higher risk for coronary heart disease (Chida & Steptoe, 2009). Such negative associations have prompted neuroimaging research into the brain correlates of high trait anger in effort to better understand its etiology and, subsequently, its possible modification or prevention.

Because of its key role in mediating defensive responses and facilitating acquisition of conditioned threat responses, the majority of existing functional neuroimaging research on trait anger has focused on the amygdala and the corticolimbic circuit (Rosell & Siever, 2016). This parallels the neuroimaging literature on disordered states characterized by high trait anger and reactive aggression, such as intermittent explosive disorder and antisocial personality disorder (McCloskey et al., 2016; Schiffer et al., 2014). For example, trait anger was positively correlated with amygdala activity to provocation in men (Repple et al., 2018), and facial expressions signaling interpersonal threat (i.e., angry faces) in men with high trait anxiety (Carré, Fisher, Manuck, & Hariri, 2012). Utilizing resting-state functional magnetic resonance imaging (rsfMRI), another study observed that trait anger was inversely correlated with the intrinsic functional connectivity between the amygdala and orbitofrontal cortex (Fulwiler, King, & Zhang, 2012). More recently, evidence has emerged for the involvement of other brain regions and circuits in the expression of trait anger, such as hyperactivity of the inferior/middle frontal gyrus, insula, and interior parietal lobule, in response to unpleasant images (Alia-Klein et al., 2018).

While these existing studies have helped outline possible brain correlates of trait anger, they have been limited in three ways. First, the majority of previous findings are derived from small sample sizes (*<*100), which is suboptimal for individual differences research using fMRI (Dubois & Adolphs, 2016). Second, as described above, prior studies have an *a priori* focus on discrete brain regions of interest (ROIs), which typically include the amygdala. Third, blood oxygen level dependent (BOLD) responses from task fMRI generally have poor psychometric properties to be considered as a reliable index of individual differences (Elliott et al., 2020).

Here, we attempted to overcome these prior limitations by conducting data-driven analyses of variability in connectome-wide functional connectivity patterns and trait anger in a large sample of young adult volunteers. First, we utilized of a dataset with substantially larger sample size (*n* = 1,048) than previous neuroimaging studies of trait anger. Second, we adopted a connectome-wide association study (CWAS) approach on our data (Shehzad et al., 2014). CWAS is particularly apt for the data-driven nature of the present investigation, as it enables the computation of multivariate connectivity patterns across the whole brain that correlates with trait anger, while making minimal assumptions about the data and not requiring *a priori* selection of networks. Third, we focused on assessing trait-like intrinsic functional architecture of brain networks, which are typically investigated using rsfMRI (Shehzad et al., 2009). Recently, we demonstrated that rsfMRI and task fMRI data could be combined to achieve more reliable estimations of intrinsic functional connectivity (GFC; general functional connectivity) by substantially increasing useable data points (Elliott et al., 2019).

Thus, through the use of CWAS, we sought to explore the neural correlates of trait anger with a reliable measure of functional connectivity across the whole brain. In order to supplement the data-driven approach, based on the existing literature emphasizing the role of the amygdala and the corticolimbic circuit (Rosell & Siever, 2016), we also aimed to test whether the functional connectivity patterns of the amygdala were associated with trait anger.

## Methods

### Participants

Neuroimaging and trait anger data were available from 1,048 young adult university students (621 women, age range 18-22 years, mean age = 19.68 years) who voluntarily participated in the Duke Neurogenetics Study (DNS) between January 2010 and November 2016. Structured clinical interview (Sheehan et al., 1998) was used to screen the participants for past or current DSM-IV (American Psychiatric Association, 1994) Axis I or select Axis II (borderline and antisocial personality) disorders. This procedure was included to facilitate comparisons with previous neuroimaging work using trait anger. The DNS was approved by the Duke University Medical Center Institutional Review Board and all participants provided written, informed consent prior to the study. To be eligible for the DNS, participants were required to be free of the following conditions: 1) medical diagnoses of cancer, stroke, head injury with loss of consciousness, untreated migraine headaches, diabetes requiring insulin treatment, chronic kidney, or liver disease; 2) use of psychotropic, glucocorticoid, or hypolipidemic medication; and 3) conditions affecting cerebral blood flow and metabolism (e.g., hypertension).

### Self-Report Measures of Trait Anger

Trait anger was assessed using the trait version of the State-Trait Anger Expression Inventory (STAXI), a 10-item self-report questionnaire designed to assess individual differences in trait anger (Spielberger, 1991). Participants were instructed to read each of the 10 statements (e.g., *When I get mad, I say nasty things*) and rate how they generally feel on a 4-point Likert-like scale from *almost never* to *almost always*. STAXI Trait anger scores ranged from 10 to 40 (mean = 15.71 ± 4.25 standard deviation).

### Image Acquisition

Each participant was scanned using one of the two identical research-dedicated GE MR750 3T scanner equipped with high-power high-duty-cycle 50-mT/m gradients at 200 T/m/s slew rate, and an eight-channel head coil for parallel imaging at high bandwidth up to 1MHz at the Duke-UNC Brain Imaging and Analysis Center. A semi-automated high-order shimming program was used to ensure global field homogeneity. A series of 34 interleaved axial functional slices aligned with the anterior commissure-posterior commissure plane were acquired for full-brain coverage using an inverse-spiral pulse sequence to reduce susceptibility artifacts (TR/TE/flip angle = 2000 ms/ 30 ms/ 60; FOV = 240 mm; 3.75 × 3.75 × 4 mm voxels; interslice skip=0). Four initial radiofrequency excitations were performed (and discarded) to achieve steady-state equilibrium. To allow for spatial registration of each participant’s data to a standard coordinate system, high-resolution three-dimensional T1-weighted structural images were obtained in 162 axial slices using a 3D Ax FSPGR BRAVO sequence (TR/TE/flip angle = 8.148 ms / 3.22 ms / 12°; voxel size = 0.9375 × 0.9375 × 1 mm; FOV = 240 mm; interslice skip=0; total scan time = 4 min and 13 s). For each participant, two back-to-back 4-minute 16-second (256 time points) rsfMRI scans were acquired. Participants were instructed to remain awake, with their eyes open during each resting-state scan. Participants also completed an emotional face-matching task (6:30 min, 195 time points), a card-guessing task (5:42 min, 171 time points), a working memory task (11:48 min, 354 time points), and a face-naming task (5:24 min, 162 time points). Detailed descriptions of these four tasks are provided in the Supplementary Materials.

### Image Processing

Anatomical images for each subject were skull stripped, intensity normalized, and nonlinearly warped to a study-specific average template in the standard stereotactic space of the Montreal Neurological Institute template using the advanced normalization tools (ANTs) SyN registration algorithm (Klein et al., 2009). Time-series images for each subject were despiked, slice time corrected, realigned to the first volume in the time-series to correct for head motion using AFNI tools (Cox, 1996), coregistered to the anatomical image using FSL’s boundary based registration (Greve & Fischl, 2009), spatially normalized into Montreal Neurological Institute space using the nonlinear ANTs SyN warp from the anatomical image, resampled to 2-mm isotropic voxels, and smoothed to minimize noise and residual difference in gyral anatomy with a Gaussian filter set at 6-mm full width at half maximum. All transformations were concatenated so that a single interpolation was performed.

Time-series images for each participant were further processed to limit the influence of motion and other artifacts. Voxelwise signal intensities were scaled to yield a time-series mean of 100 for each voxel. Motion regressors were created using each subject’s six motion correction parameters (3 rotation and 3 translation) and their first derivatives (Satterthwaite et al., 2013) yielding 12 motion regressors. White matter and cerebrospinal fluid nuisance regressors were created using CompCor (Behzadi, Restom, Liau, & Liu, 2007). Images were bandpass filtered to retain frequencies between 0.008 and 0.1 Hz, and volumes exceeding 0.25-mm framewise displacement or 1.55 standardized DVARS (Power, Mitra, Laumann, Snyder, Schlaggar, & Petersen, 2014) were censored. Nuisance regression, bandpass filtering, and censoring for each time-series was performed in a single processing step using AFNI’s *3dTproject*.

### General Functional Connectivity

In order to compute GFC, identical time-series preprocessing steps were applied to both rsfMRI and task fMRI data with one exception. As task fMRI data include task-evoked coactivation that may drive functional connectivity, signal due to task structure was added as an additional nuisance covariate and removed from the time-series (Fair et al., 2007). This approach has been shown to improve the ability of the functional connectivity estimates in predicting behavioral measures (Elliott et al., 2019). GFC was computed by calculating functional connectivity after concatenating all rsfMRI and task fMRI scans to form a single time-series. All 1,048 participants had more than or equal to 185 time points left after censoring.

### Connectome-wide Association Study

Following the preprocessing steps that included censoring and nuisance regressions to limit the influence of head motion and other artifacts, fMRI time-series data were extracted from the Power 264 atlas, which parcellated the brain into 264 regions (Power et al., 2011). BOLD data were averaged within 5 mm spheres surrounding each 264 coordinates in the parcellation. As noted elsewhere (Elliott et al., 2019), average time-series data were extracted independently from each scan session. This allowed the time-series from rsfMRI and task fMRI data to be flexibly concatenated and recombined. Extracted time-series data for each participant were then processed using a CWAS approach (Shehzad et al., 2014). CWAS utilizes multi-dimensional matrix regression (MDMR) to identify seed regions with whole-brain patterns of intrinsic functional connectivity that are associated with another variable. In brief, CWAS can be summarized into three steps. First, seed-based functional connectivity analysis is performed with a single region of interest (ROI) to generate a whole-brain functional connectivity map for each participant. Second, the average distance (1-Pearson correlation *r*) between each pair of participant’s functional connectivity maps is computed, resulting in a distance matrix encoding the multivariate similarity between each participant’s connectivity map. Finally, MDMR is used to generate a pseudo-*F* statistic quantifying the strength of the association between the variable of interest (e.g., trait anger) and the distance matrix created in the second step. These three steps are repeated for each of the 264 ROIs, resulting in a whole-brain map that represents the association between trait anger and whole-brain connectivity at each ROI. Age and sex were included as covariates, and 100,000 permutations were performed to generate *p*-values. To account for false positives across the 264 ROIs, a false discovery rate (FDR) correction was applied (Benjamini & Hochberg, 1995). Statistical significance threshold was set at *q* = 0.05.

### Seed-based Analyses

MDMR identifies a set of ROIs with patterns of whole-brain connectivity associated with a variable of interest. However, the nature in which the connectivity of these MDMR-selected ROIs relates to said variable remains unclear. To address this, previous research using CWAS (Elliott et al., 2018; Shehzad et al., 2014) has utilized traditional seed-based connectivity follow-up analyses to better understand the networks and brain regions that drive the associations discovered through MDMR. We followed the same procedure described in our previous work (Elliott et al., 2018). Namely, seed-based connectivity maps were created and correlation coefficients at each voxel were converted to *z* statistics via the Fisher’s *r*-to-*z* transform. Then, correlations between these connectivity values and trait anger were calculated across the whole brain, with age and sex included as covariates. Importantly, these follow-up analyses do not represent independent statistical tests because they were performed *post hoc* to the family-wise error-controlled MDMR findings. Accordingly, these follow-up analyses were not thresholded to visualize all information that was relevant to the MDMR step. Finally, to characterize the overall MDMR seed-based functional connectivity patterns, functional connectivity estimates were summarized for each of seven previously identified canonical intrinsic networks (Yeo et al., 2011) to quantify the relative contributions of each network to the observed associations with trait anger.

Finally, seed-based analyses using *a priori* amygdala ROIs were performed, as the Power 264 atlas used in MDMR analyses does not cover the amygdala. Amygdala ROIs were defined using a high-resolution template generated from the 168 Human Connectome Project dataset (Tyszka & Pauli, 2016). For completeness, whole-brain functional connectivity analyses were performed separately for the basolateral (BL) and centromedial (CM) amygdala subregions on each hemisphere, yielding four amygdala ROIs. For this step, we followed the same procedures described above with the exception of implementing statistical thresholds, as these analyses do represent independent statistical tests from MDMR. Nonparametric permutation tests (*n* = 5,000) were performed on the data to determine significant voxels at *p* < 0.05 corrected for multiple comparisons, using the *randomise* tool along with the threshold-free cluster enhancement method implemented in FSL (Smith and Nichols, 2009; Winkler et al., 2014).

## Results

### Multidimension Matrix Regression

MDMR identified three brain regions for which whole-brain functional connectivity patterns significantly varied as a function of trait anger (Figure 1). These regions were the left supplementary motor area (MNI-3, 2, 53; FDR-corrected *p* = 0.005), right supplementary motor area (MNI 7, 8, 51; FDR-corrected *p* = 0.008), and right lateral frontal pole (MNI 49, 35, −12; FDR-corrected *p* < 0.001).

**Figure 1.**
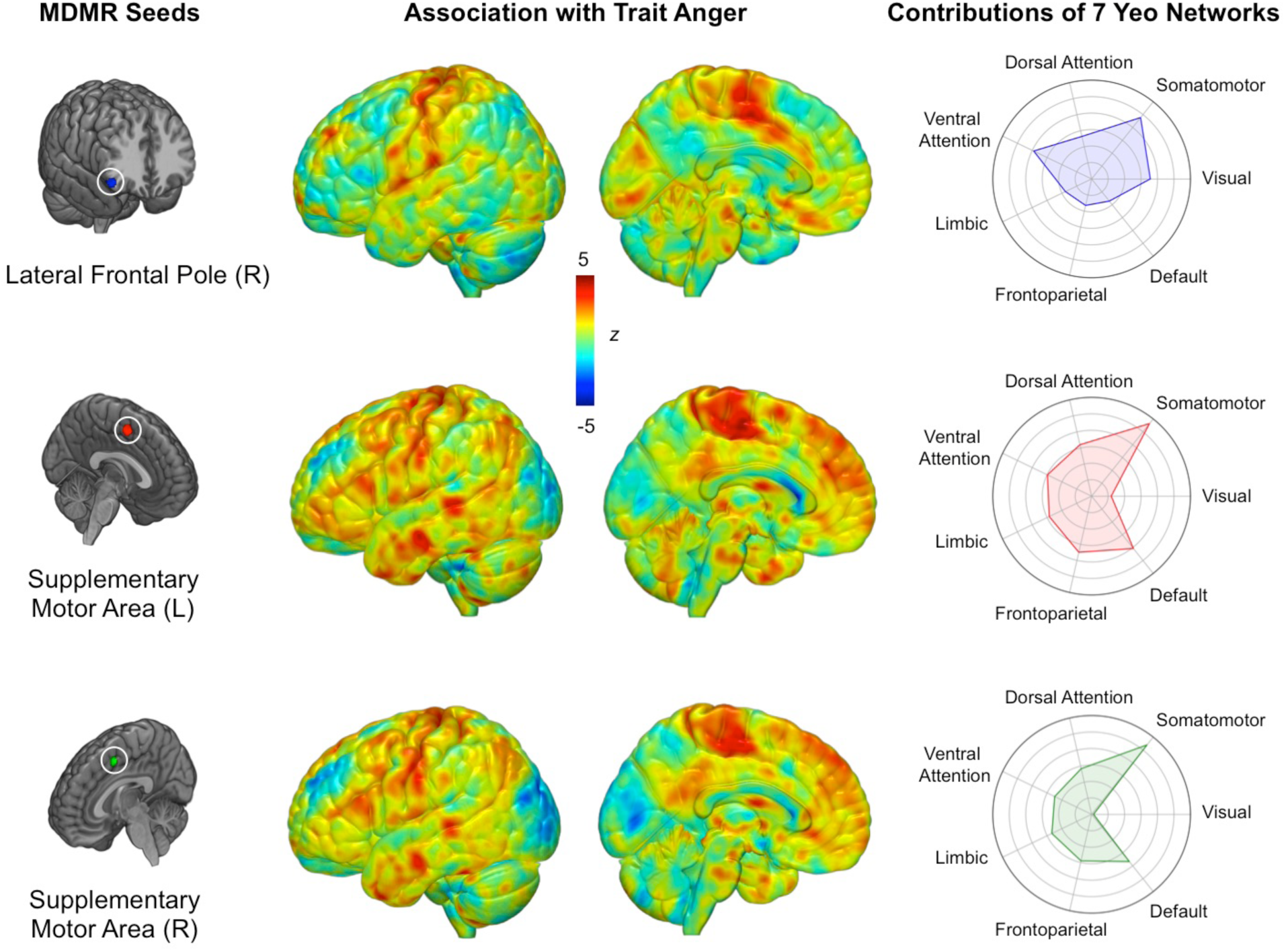
Data-driven MDMR analysis identified three regions with whole-brain functional connectivity patterns significantly associated with trait anger (left panel). Connectome-wide functional connectivity patterns for each MDMR-derived seed region as a function of trait anger (middle panel). Relative involvement of seven canonical functional networks to each connectivity pattern (right panel). Values farther toward the outer circle indicate greater functional connectivity to the corresponding network in high trait anger.

### Follow-up Functional Connectivity Analyses

Follow-up seed-based connectivity analyses revealed that while showing some variability, the three MDMR-selected ROIs exhibited largely overlapping patterns of whole-brain functional connectivity as a function of trait anger that converged on several brain regions (Figure 1). When mean functional connectivity estimates were calculated for each of seven canonical intrinsic networks (Yeo et al., 2011), high trait anger was associated with increased functional connectivity between all three MDMR-derived ROIs and the somatomotor network (SMN). Surveying each of the three MDMR-derived ROIs separately, the left and right supplementary motor areas showed nearly identical functional connectivity patterns: higher trait anger was associated with 1) increased functional connectivity with the default mode network (DMN) and 2) decreased functional connectivity with the visual network (VN). Conversely, the functional connectivity of the right lateral frontal pole did not exhibit such associations with either the DMN or the VN.

### Amygdala Functional Connectivity Analyses

Of the four amygdala ROIs, the left CM did not yield significant associations. The remaining three amygdala ROIs exhibited variability in functional connectivity associated with trait anger across a network of cortical and subcortical brain regions. Converging results were observed across these three seed ROIs with minor differences (Figure 2). Generally, higher trait anger was associated with increased connectivity between these ROIs and the dorsomedial prefrontal cortex (dmPFC), dorsal anterior cingulate cortex (dACC), thalamus, caudate, putamen, nucleus accumbens, supplementary motor area (SMA), medial frontal pole (FPm), middle frontal gyrus (MFG), temporal pole (TP), and brainstem (see Supplementary Materials for additional details). There were no brain regions exhibiting decreased functional connectivity with any amygdala ROIs as a function of trait anger.

**Figure 2.**
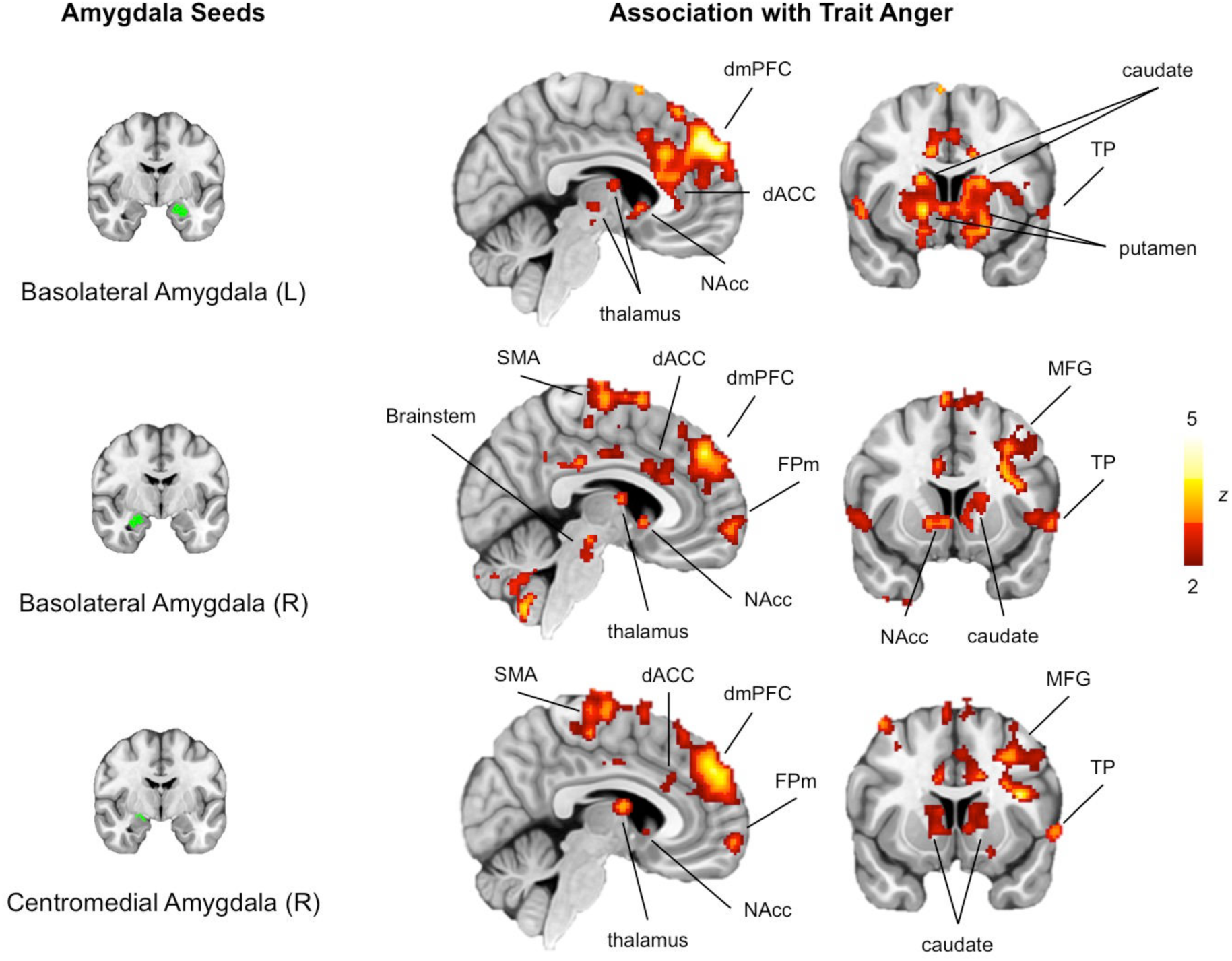
Seed-based analysis using amygdala ROIs (left panel) showed that high trait anger was associated with hyperconnectivity of the amygdala with a network of cortical and subcortical brain regions (right panel). Results shown here are visualized with a statistical threshold of *p* < 0.05, corrected for multiple comparisons across the whole brain. No brain regions survived this threshold when the left centromedial amygdala was used as a seed region. *Note:* dACC (dorsal anterior cingulate cortex), dmPFC (dorsomedial prefrontal cortex), FPm (medial frontal pole), NAcc (nucleus accumbens), MFG (middle frontal gyrus), SMA (supplementary motor area), TP (temporal pole)

## Discussion

By utilizing a data-driven approach on a reliable measure of the functional connectome in a large sample of 1,048 young adult volunteers, we provide novel evidence that trait anger is associated with variability in the functional connectivity patterns of multiple networks supporting somatomotor, affective, self-referential, and visual information processes. The connectome-wide data-driven MDMR analysis identified three brain regions, left and right supplementary motor area and right lateral frontal pole, with whole-brain functional connectivity patterns significantly associated with trait anger. Higher trait anger was associated with hyperconnectivity between all three of these regions and the somatomotor network. Further analyses revealed that, as a function of trait anger, the bilateral supplementary motor area but not the right lateral frontal pole showed hyperconnectivity with the default mode network and hypoconnectivity with the visual network. These findings expand upon previous work focused on associations between trait anger and variability within a corticolimbic circuit supporting threat learning to include action and action-planning brain regions and networks.

Specifically, all three brain regions identified through the MDMR analysis contribute to action and action-planning. The supplementary motor area is a well-known component of the motor cortex, functioning to prepare for voluntary action or movement (Cunnington et al., 2003). The lateral frontal pole is associated with the cognitive control of behavior (Orr, Smolker & Banich, 2015), which includes, of particular relevance to the current study, socioemotional approach-avoidance actions (Bramson et al., 2020). These findings are consistent with previously documented action-related aspects of trait anger, such as increased approach motivation (Harmon-Jones, 2003) and poor inhibitory control (Wilkowski & Robinson, 2008).

The involvement of action-related networks in the expression of trait anger was further reflected in the specific patterns of functional connectivity displayed by these MDMR-derived brain regions. The somatomotor network, which encompasses brain regions including the primary motor cortex (Brodmann area 4), premotor area (area 6), primary somatosensory cortex (areas 1, 2, 3), early somatosensory area (area 5L), and parts of the midcingulate sulcus, showed hyperconnectivity with all three seeds as a function of trait anger. One possible interpretation of this pattern associated with higher trait anger is that it may reflect a lowered threshold for action based on somatosensory information, as the three action-related brain regions became more synchronized with the SMN as a function of trait anger. Such speculation offers an interesting neural basis to an important characteristic of trait anger – that it reflects a greater propensity to provoked action.

Some diverging patterns of connectivity across the three MDMR-derived brain regions were observed as well. The default mode network, best known for its integral role in self-projection and mental simulations (Buckner & DiNicola, 2019), showed hyperconnectivity with the bilateral supplementary motor area as a function of trait anger, while an opposite pattern was found for the visual network. Recently, it was reported that major functional networks form a distinct spatial organization that follows a macroscale gradient (Buckner & DiNicola, 2019; Margulies et al., 2016). Interestingly, the DMN is located at one extreme end of this gradient, maximally separated from both SMN and VN, which are situated at the opposite end (Margulies et al., 2016). This spatial gradient also reflects the orientation of the functional processes associated with each network, such that the DMN reflects processes oriented towards the self, whereas SMN and VN represent processes oriented towards the environment. From this perspective, trait anger may reflect the psychological manifestation of the changes in the distance between an internally oriented brain process (i.e., DMN activity) and externally oriented brain processes (i.e.. SMN, VN activity) along this gradient. In other words, trait anger could be thought of as action-readiness (supplementary motor area) becoming more influenced by self-referential and somatomotor information (hyperconnectivity with the DMN, SMN) while relying less on visual information (hypoconnectivity with the VN).

Our data-driven connectome-wide MDMR analysis was augmented by a seed-based analysis using the amygdala, which revealed a broad network of cortical and subcortical brain regions with altered functional connectivity as a function of trait anger. Previous investigations on the neural correlates of trait anger have typically focused on the amygdala (Repple et al., 2018; Wang et al., 2017; Carré et al., 2012; Carlson, Greenberg & Mujica-Parodi, 2010) and its connectivity with the prefrontal cortices (Beyer et al., 2014; Fulwiler et al., 2012) based on *a priori* theoretical and empirical work (LeDoux, 1996; Marsh & Blair, 2008). While other studies have individually reported a number of brain regions beyond the amygdala associated with trait anger, such as the thalamus (Alia-Klein et al., 2018; Herpertz et al., 2017), striatum (da Cunha-Bang et al., 2017), and anterior cingulate cortex (Repple et al., 2018), our analysis demonstrates that all of these neural sites reflect individual differences in trait anger with regards to the amygdala. Above and beyond the methodological differences (e.g., task vs. resting fMRI), it is likely that the large sample size we have employed for the present study secured sufficient statistical power to observe these associations that were previously undetected. Of particular interest from this final set of analyses is the hyperconnectivity between the amygdala and the dmPFC in high trait anger. While a direct evidence for trait anger is sparse, hyperconnectivity of the amygdala-dmPFC circuit is often reported to be associated with negative psychological and physical health outcomes, such as heightened and sustained anxiety (Kim et al., 2011; Vytal et al., 2014), as well as enhanced inflammatory responses to stress (Muscatell et al., 2015). Perhaps the amygdala-dmPFC hyperconnectivity observed in individuals with high trait anger may reflect their relative susceptibility to negative health outcomes.

The present study is not without limitations that can be addressed in future research. First, as the DNS sampled high functioning university students, the generalizability of the present findings to a broader population is yet to be confirmed. Future studies utilizing large-scale, population-representative datasets would be able to address this issue. Second, the assessment of trait anger relied on self-report, via a standardized questionnaire. Development of a more objective measurement of trait anger (e.g., behavioral observation) would be helpful in overcoming the limitations associated with self-report. Third, the current findings are derived from generally low levels of self-reported trait anger. Thus, we caution against making broader inferences from our data (e.g., to disordered states such as the intermittent explosive disorder). Finally, we did not observe strong evidence for the involvement of the limbic network with regards to the three MDMR-derived seed regions and trait anger. While this may be the case, it is noteworthy that out of the seven canonical functional networks, the limbic network displayed the lowest level of reliability in other large-scale fMRI datasets (Elliott et al., 2019). As such, we cannot rule of the possibility that our observations in the present study may have been affected by differences in reliability across the limbic network vs. other canonical functional networks.

These limitations notwithstanding, our study design and analyses allowed us to address key limitations of prior studies, which includes small sample size, reliance on *a priori* ROIs, and poor psychometric properties of BOLD responses in task fMRI. Our large sample size enabled us to employ data-driven, exploratory approaches on the whole brain functional networks. The functional connectome was generated from functional connectivity data that was shown to have good reliability and predictive utility, both of which are necessary psychometric properties for fMRI-based individual differences research (Elliott et al., 2020).

To summarize, the present study leveraged a large dataset and an unconstrained connectome-wide MDMR approach to highlight a collection of brain regions and canonical intrinsic networks associated with trait anger, many of which were previously undetected. Findings from our data-driven, exploratory analyses could help generate new hypotheses for future neuroimaging research of trait anger, as well as related constructs. These include emotional lability and irritability, both of which have implications for adversely affecting the developing brain and mental health (Bennett, Somandepalli, Roy & Di Martino, 2017; Dennis, Humphreys, King, Thompson & Gotlib, 2019). Future studies could be designed to utilize the present findings and examine the intrinsic functional networks in relevant psychiatric disorders such as the intermittent explosive disorder (Gan et al., 2019), in order to elucidate the utility of trait anger as a potential transdiagnostic marker.

## References

Alia-Klein, N., Preston-Campbell, R. N., Moeller, S. J., Parvaz, M. A., Bachi, K., Gan, G.,… Goldstein, R. Z. (2018). Trait anger modulates neural activity in the fronto-parietal attention network. PLoS One, 13, e0194444. http://dx.doi.org/10.1371/journal.pone.0194444

American Psychiatric Association. (1994). Diagnostic and Statistical Manual of Mental Disorders (4th ed.). Washington, DC: Author

Behzadi, Y., Restom, K., Liau, J., & Liu, T. T. (2007). A component based noise correction method (CompCor) for BOLD and perfusion based fMRI. Neuroimage, 37, 90–101. http://dx.doi.org/10.1016/j.neuroimage.2007.04.042

Benjamini, Y., & Hochberg, Y. (1995). Controlling the false discovery rate: a practical and powerful approach to multiple testing. Journal of the Royal Statistical Society. Series B (Methodological), 57, 289–300. http://dx.doi.org/10.1111/j.2517-6161.1995.tb02031.x

Bennett, R. H., Somandepalli, K., Roy, A. K., & Di Martino, A. (2017). The neural correlates of emotional lability in children with autism spectrum disorder. Brain Connectivity, 7, 281–288. http://dx.doi.org/10.1089/brain.2016.0472

Bettencourt, B. A., Talley, A., Benjamin, A. J., & Valentine, J. (2006). Personality and aggressive behavior under provoking and neutral conditions: A meta-analytic review. Psychological Bulletin, 132, 751– 777. http://dx.doi.org/10.1037/0033-2909.132.5.751

Beyer, F., Münte, T. F., Wiechert, J., Heldmann, M., & Krämer, U. M. (2014). Trait aggressiveness is not related to structural connectivity between orbitofrontal cortex and amygdala. PLoS One, 9, e101105. http://dx.doi.org/10.1371/journal.pone.0101105

Bramson, B., Folloni, D., Verhagen, L., Hartogsveld, B., Mars, R. B., Toni, I.,… & Roelofs, K. (2020). Human lateral frontal pole contributes to control over emotional approach– avoidance actions. Journal of Neuroscience, 40, 2925–2934. http://dx.doi.org/10.1523/JNEUROSCI.2048-19.2020

Buckner, R. L., & DiNicola, L. M. (2019). The brain’s default network: updated anatomy, physiology and evolving insights. Nature Reviews Neuroscience, 20, 593–608. http://dx.doi.org/10.1038/s41583-019-0212-7

Carlson, J. M., Greenberg, T., & Mujica-Parodi, L. R. (2010). Blind rage? Heightened anger is associated with altered amygdala responses to masked and unmasked fearful faces. Psychiatry Research: Neuroimaging, 182, 281–283. http://dx.doi.org/10.1016/j.pscychresns.2010.02.001

Carré, J. M., Fisher, P. M., Manuck, S. B., & Hariri, A. R. (2012). Interaction between trait anxiety and trait anger predict amygdala reactivity to angry facial expressions in men but not women. Social Cognitive and Affective Neuroscience, 7, 213–221. http://dx.doi.org/10.1093/scan/nsq101

Chida, Y., & Steptoe, A. (2009). The association of anger and hostility with future coronary heart disease: A meta-analytic review of prospective evidence. Journal of the American College of Cardiology, 53, 936–946. http://dx.doi.org/10.1016/j.jacc.2008.11.044

Cox R. W. (1996). AFNI: software for analysis and visualization of functional magnetic resonance neuroimages. Computational Biomedical Research, 29, 162–173. http://dx.doi.org/10.1006/cbmr.1996.0014

Cunnington, R., Windischberger, C., Deecke, L., & Moser, E. (2003). The preparation and readiness for voluntary movement: a high-field event-related fMRI study of the Bereitschafts-BOLD response. Neuroimage, 20, 404–412. http://dx.doi.org/10.1016/s1053-8119(03)00291-x

da Cunha-Bang, S., Fisher, P. M., Hjordt, L. V., Perfalk, E., Persson Skibsted, A., Bock, C.,… & Knudsen, G. M. (2017). Violent offenders respond to provocations with high amygdala and striatal reactivity. Social Cognitive and Affective Neuroscience, 12, 802–810. http://dx.doi.org/10.1093/scan/nsx006

Dennis, E. L., Humphreys, K. L., King, L. S., Thompson, P. M., & Gotlib, I. H. (2019). Irritability and brain volume in adolescents: cross-sectional and longitudinal associations. Social Cognitive and Affective Neuroscience, 14, 687–698. http://dx.doi.org/10.1093/scan/nsz053

Dubois, J., & Adolphs, R. (2016). Building a science of individual differences from fMRI. Trends in Cognitive Sciences, 20, 425–443. http://dx.doi.org/ doi:10.1016/j.tics.2016.03.014

Eckhardt, C. I., Jamison, T. R., & Watts, K. (2002). Anger experience and expression among male dating violence perpetrators during anger arousal. Journal of Interpersonal Violence, 17, 1102–1114. http://dx.doi.org/10.1177/088626002236662

Elliott, M. L., Knodt, A. R., Cooke, M., Kim, M. J., Melzer, T., Keenan, R.,… & Hariri, A. R. (2019). General Functional Connectivity: shared features of resting-state and task fMRI drive reliable and heritable individual differences in functional brain networks. Neuroimage, 189, 516–532. doi:10.1016/j.neuroimage.2019.01.068

Elliott, M. L., Knodt, A. R., Ireland, D., Morris, M. L., Poulton, R., Ramrakha, S.,… & Hariri, A. R. (2020). What is the test-retest reliability of common task-functional MRI measures? New empirical evidence and a meta-analysis. Psychological Science, 31, 792–806.

Elliott, M. L., Romer, A., Knodt, A. R., & Hariri, A. R. (2018). A connectome-wide functional signature of transdiagnostic risk for mental illness. Biological Psychiatry, 84, 452–459. http://dx.doi.org/10.1016/j.biopsych.2018.03.012

Fair, D. A., Schlaggar, B. L., Cohen, A. L., Miezin, F. M., Dosenbach, N. U. F., Wenger, K. K.,… & Petersen, S. E. (2007). A method for using blocked and event-related fMRI data to study “resting state” functional connectivity. Neuroimage, 35, 394–405. http://dx.doi.org/10.1016/j.neuroimage.2006.11.051

Fulwiler, C. E., King, J. A., & Zhang, N. (2012). Amygdala-orbitofrontal resting-state functional connectivity is associated with trait anger. Neuroreport, 23, 606–610. http://dx.doi.org/10.1097/WNR.0b013e3283551cfc

Gan, G., Zilverstand, A., Parvaz, M. A., Preston-Campbell, R. N., d’Oleire Uquillas, F., Moeller, R. J.,… Alia-Klein, N. (2019). Habenula-prefrontal resting-state connectivity in reactive aggressive men – a pilot study. Neuropharmacology, 156, 107396. http://dx.doi.org/10.1016/j.neuropharm.2018.10.025

Greve, D. N., & Fischl, B. (2009). Accurate and robust brain image alighnment using boundary-based registration. Neuroimage, 48, 63–72. http://dx.doi.org/10.1016/j.neuroimage.2009.06.060

Harmon-Jones, E. (2003). Anger and the behavioural approach system. Personality and Individual Differences, 35, 995–1005. http://dx.doi.org/10.1016/S0191-8869(02)00313-6

Herpertz, S. C., Nagy, K., Ueltzhöffer, K., Schimitt, R., Mancke, F., Schmahl, C.,… & Berstch, K. (2017). Brain mechanisms underlying reactive aggression in borderline personality disorder-sex matters. Biological Psychiatry, 82, 257–266. http://dx.doi.org/10.1016/j.biopsych.2017.02.1175

Kim, M. J., Gee, D. G., Loucks, R. A., Davis, F. C., & Whalen, P. J. (2011). Anxiety dissociates dorsal and ventral medial prefrontal cortex functional connectivity with the amygdala at rest. Cerebral Cortex, 21, 1667–1673. http://dx.doi.org/10.1093/cercor/bhq237

Klein, A., Andersson, J., Ardekani, B. A., Ashburner, J., Avants, B., Chiang, M. C.,… Parsey, R. V. (2009). Evaluation of 14 nonlinear deformation algorithms applied to human brain MRI registration. Neuroimage, 46, 786–802. http://dx.doi.org/10.1016/j.neuroimage.2008.12.037

LeDoux, J. E. (1996). The Emotional Brain. New York, NY: Simon and Shuster.

Margulies, D. S., Ghosh, S. S., Goulas, A., Falkiewicz, M., Huntenberg, J. M., Langs, G.,… Smallwood, J. (2016). Situating the default-mode network along a principal gradient of macroscale cortical organization. Proccedings of the National Academy of Sciences U.S.A., 113, 12574–12579. http://dx.doi.org/10.1073/pnas.1608282113

Marsh, A. A., & Blair, R. J. (2008). Deficits in facial affect recognition among antisocial populations: a meta-analysis. Neuroscience and Biobehavioral Reviews, 32, 454–465. http://dx.doi.org/10.1016/j.neubiorev.2007.08.003

McCloskey, M. S., Phan, K. L., Angstadt, M., Fettich, K. C., Keedy, S., & Coccaro, E. F. (2016). Amygdala hyperactivation to angry faces in intermittent explosive disorder. Journal of Psychiatric Research, 79, 34–41. http://dx.doi.org/10.1016/j.jpsychires.2016.04.006

Muscatell, K. A., Dedovic, K., Slavich, G. M., Jarcho, M. R., Breen, E. C., Bower, J. E.,… & Eisenberger, N. I. (2015). Greater amygdala activity and dorsomedial prefrontal–amygdala coupling are associated with enhanced inflammatory responses to stress. *Brain*, Behavior, and Immunity, 43, 46–53. http://dx.doi.org/10.1016/j.bbi.2014.06.201

Orr, J. M., Smolker, H. R., & Banich, M. T. (2015). Organization of the human frontal pole revealed by large-scale DTI-based connectivity: Implications for control of behavior. PLoS ONE, 10, e0124797. http://dx.doi.org/10.1371/journal.pone.0124797

Power, J. D., Cohen, A. L., Nelson, S. M., Wig, G. S., Barnes, K. A., Church, J. A., Vogel, A. C., Laumann, T. O., Miezin, F. M., Schlaggar, B. L., & Petersen, S. E. (2011). Functional network organization of the human brain, Neuron, 72, 665–678. https://dx.doi.org/10.1016/j.neuron.2011.09.006

Power, J. D., Mitra, A., Laumann, T. O., Snyder, A. Z., Schlaggar, B. L., & Petersen, S. E. (2014). Methods to detect, characterize, and remove motion artifact in resting state fMRI. Neuroimage, 84, 320–341. http://dx.doi.org/10.1016/j.neuroimage.2013.08.048

Repple, J., Habel, U., Wagels, L., Pawliczek, C. M., Schneider, F., & Kohn, N. (2018). Sex differences in the neural corrleates of aggression. Brain Structure and Function, 223, 4415–4124. http://dx.doi.org/10.1007/s00429-018-1739-5

Rosell, D. R., & Siever, L. J. (2016). The neurobiology of aggression and violence. CNS Spectrums, 20, 254–279. http://dx.doi.org/10.1017/S109285291500019X

Satterthwaite, T. D., Elliott, M. A., Gerraty, R. T., Ruparel, K., Loughead, J., Calkins, M. E.,… Wolf, D. H. (2013). An improved framework for confound regression and filtering for control of motion artifact in the preprocessing of resting-state functional connectivity data. Neuroimage, 64, 240–256. http://dx.doi.org/10.1016/j.neuroimage.2012.08.052

Schiffer, B., Pawliczek, C., Müller, B., Forsting, M., Gizewski, E., Leygraf, N.,… & Hodgins, S. (2014). Neural mechanisms underlying cognitive control of men with lifelong antisocial behavior. Psychiatry Research: Neuroimaging, 222, 43–51. http://dx.doi.org/10.1016/j.pscychresns.2014.01.008

Sheehan, D. V., Lecrubier, Y., Sheehan, H. K., Amorim, P., Janavs, J., Weiller, E.,… Dunbar, G. C. (1998). The Mini-International Neuropsychiatric Interview (M.I.N.I.): The development and validation of a Structured Diagnostic Psychiatric Interview for DSM–IV and ICD-10. Journal of Clinical Psychiatry, 59, 22–33.

Shehzad, Z., Kelly, C., Reiss, P. T., Cameron Craddock, R., Emerson, J. W., McMahon, K.,… Milham, M. P. (2014). A multivariate distance-based analytic framework for connectome-wide association studies. Neuroimage, 93, 74–94. http://dx.doi.org/10.1016/j.neuroimage.2014.02.024

Shehzad, Z., Kelly, A. M. C., Reiss, P. T., Gee, D. G., Gotimer, K., Uddin, L. Q.,… Milham, M. P. (2009): The resting brain: Unconstrained yet reliable. Cereb Cortex 19, 2209–2229. http://dx.doi.org/0.1093/cercor/bhn256

Smith, S. M., & Nichols, T. E. (2009). Threshold-free cluster enhancement: addressing problems of smoothing, threshold dependence and localization in cluster inference. Neuroimage, 44, 83–98. http://dx.doi.org/10.1016/j.neuroimage.2008.03.061

Spielberger, C. D. (1991). State-Trait Anger Expression Inventory. Odessa, FL: Psychological Assessment Resources Inc.

Tyszka, J. M., & Pauli, W. M. (2016). *In vivo* delineation of subdivisions of the human amygdaloid complex in a high-resolution group template. Human Brain Mapping, 37, 3979–3998. http://dx.doi.org/10.1002/hbm.23289

Wang, Y., Kong, F., Kong, X., Zhao, Y., Lin, D., & Liu, J. (2017). Unsatisfied relatedness, not competence or autonomy, increases trait anger through the right amygdala. *Cognitive*, Affective, and Behavioral Neuroscience, 17, 932–938. http://dx.doi.org/10.3758/s13415-017-0523-y.

Wilkowski, B. M., & Robinson, M. D. (2008). The cognitive basis of trait anger and reactive aggression: An integrative analysis. Personality and Social Psychology Review, 12, 3–21. http://dx.doi.org/10.1177/1088868307309874

Winkler, A. M., Ridgway, G. R., Webster, M. A., Smith, S. M., & Nichols, T. E. (2014). Permutation inference for the general linear model. Neuroimage, 92, 381–397. http://dx.doi.org/10.1016/j.neuroimage.2014.01.060

Vytal, K. E., Overstreet, C., Charney, D. R., Robinson, O. J., & Grillon, C. (2014). Sustained anxiety increases amygdala–dorsomedial prefrontal coupling: a mechanism for maintaining an anxious state in healthy adults. Journal of Psychiatry & Neuroscience, 39, 321–329.

Yeo, B. T., Krienen, F. M., Sepulcre, J., Sabuncu, M. R., Lashkari, D., Hollinshead, M.,… Buckner, R. L. (2011). The organization of the human cerebral cortex estimated by intrinsic functional connectivity. Journal of Neurophysiology, 106, 1125–1165. http://dx.doi.org/10.1152/jn.000338.2011

