## Supplementary Materials for "A Connectome-wide Functional Signature of Trait Anger"

### Supplementary Methods

Summarized below are descriptions of the four experimental paradigms used for the task fMRI scans. We note that these descriptions are also available in our previous published work (Farber et al., 2019; Forbes et al., 2009; Kim et al., 2018).

### Emotional Face-matching Task

The experimental fMRI paradigm consists of four blocks of a perceptual face-matching task interleaved with five blocks of a sensorimotor control task. The DNS version of this paradigm consists of one block each of fearful, angry, surprised, and neutral facial expressions presented in a pseudorandom order across participants. During face-matching blocks, participants view a trio of faces and select one of two faces (on the bottom) identical to a target face (on the top). Each face processing block consists of six images, balanced for gender, all of which were derived from a standard set of pictures of facial affect (Ekman & Friesen, 1976). During the sensorimotor control blocks, participants view a trio of simple geometric shapes (circles and vertical and horizontal ellipses) and select one of two shapes (bottom) that are identical to a target shape (top). Each sensorimotor control block consists of six different shape trios. All blocks are preceded by a brief instruction ("Match Faces" or "Match Shapes") that lasts 2 s. In the task blocks, each of the six face trios is presented for 4 s with a variable interstimulus interval (ISI) of 2-6 s (mean = 4 s) for a total block length of 48 s. A variable ISI is used to minimize expectancy effects and resulting habituation, and maximize amygdala reactivity throughout the paradigm. In the control blocks, each of the six shape trios is presented for 4 s with a fixed ISI of 2 s for a total block length of 36 s. Total task time is 390 s.

**Card-guessing Task**

Our blocked-design number-guessing paradigm consists of a pseudorandom presentation of three blocks of predominantly positive feedback (80% correct guess), three blocks of predominantly negative feedback (20% correct guess) and three control blocks. There are five trials in 3 seconds to guess, via button press, whether the value of a visually presented card is lower or higher than 5 (index and middle finger, respectively). The numerical value of the card is then presented for 500 milliseconds and followed by appropriate feedback (green upward-facing arrow for positive feedback; red downward-facing arrow for negative feedback) for an additional 500 milliseconds. A crosshair is then presented for 3 seconds, for a total trial length of 7 seconds. Each block comprises five trails, with three blocks each of predominantly positive feedback (80% correct) and three of predominantly negative feedback (20% correct) interleaved with three control blocks. During control blocks, participants are instructed to simply make button presses during the presentation of an "x" (3 seconds), which is followed by an asterisk (500 milliseconds) and a yellow circle (500 milliseconds). Each block is preceded by an instruction of "Guess Number" (positive or negative feedback blocks) or "Press Button" (control blocks) for 2 seconds resulting in a total block length of 38 seconds and a total task length of 342 seconds. Participants were unaware of the fixed outcome probabilities associated with each block and were led to believe that their performance would determine a net monetary gain at the end of the scanning session. Instead, all participants received $10. We included one incongruent trial within each task block (e.g., one of five trials during positive feedback blocks was incorrect resulting in negative feedback) to prevent participants from anticipating the feedback for each trial and to maintain participants' engagement and motivation to perform well.

**Working Memory Task**

An event-related working memory paradigm adapted from Tan et al. (2007). The paradigm included 10 trials for each of 6 different conditions, including 3 control conditions, consisting only of a 3s response phase, and 3 working memory (WM) conditions, consisting of a 0.5s encoding phase followed by a 4s maintenance interval and a 3s response phase. Control and WM conditions were interleaved with jittered rest intervals lasting 4s to 8.5s for a total scan length of 11m 48s. Responses were recorded via an MR-compatible button box using the index (left button) and middle (right button) fingers of the dominant hand. During the control conditions, participants performed 1) a simple motor task (M) in which they pressed either the left or the right button according to a prompt, 2) a numerical size judgment task (J) in which they chose the number on the left or right based on an instruction to choose either the larger or the smaller number, and 3) a numerical computation and size judgment task (CJ) in which they performed a numerical subtraction of 2 or 3 from either the left or right number, and made a numerical size judgment as instructed. In the first WM condition, participants viewed 2 numbers during the brief encoding phase, then recalled the numbers and performed a numerical size judgment as instructed (E_RJ). In the second WM condition, the participants additionally performed subtraction of 2 or 3 from one of the remembered numbers as indicated before making the numerical size judgment during recall (E_RCJ). In the final WM condition, participants performed subtraction of 2 or 3 from one of the 2 numbers during the brief encoding phase, then recalled the resulting two numbers and performed a numerical size judgment as instructed during the response phase after the maintenance interval (EC_RJ). In each WM condition trial, all the numbers were single digits from 0 to 9; the two numbers on which the numerical size judgment was ultimately performed (after numerical computation if applicable) were equally balanced across 0 to 9, and equally likely to differ by either 1 or 3 units. Numerical computation was equally likely on the left or right number, with correct responses equally balanced on the left or right, and equally likely to be the larger or smaller number for each WM trial type. The trials were performed in an order that was optimized using a sequencing program (Wager & Nichols, 2003).

**Face-naming Task**

Our fMRI paradigm consists of the encoding and subsequent recall of novel face-name pairs (Zeineh et al., 2003). A distractor task (odd/even number identification) is interleaved between encoding and recall blocks to prevent maintenance of information in working memory. During each of four encoding blocks, subjects view six novel face-name pairs for 3.5 seconds each. During each of four recall blocks, subjects view six faces each presented for 2 seconds and immediately followed by an incomplete name fragment for 1 second during which they are required by forced-choice to determine if the fragment is correct or incorrect. A 1 second inter-trial interval is used during recall blocks. During each of four distractor blocks, subjects view six different numbers for 3.5 seconds each and are required to determine if the numbers are odd or even. Total task length is 324 seconds.

**References**

Ekman, P., & Friesen, W. (1976). *Pictures of Facial Affect*. Palo Alto, CA: Consulting Psychologists Press.

Farber, M. J., Kim, M. J., Knodt, A. R., & Hariri, A. R. (2019). Maternal overprotection in childhood is associated with amygdala reactivity and structural connectivity in adulthood. *Developmental Cognitive Neuroscience, 40*, 100711.

Forbes, E. E., Brown, S. M., Kimak, M., Ferrell, R. E., Manuck, S. B., & Hariri, A. R. (2009). Genetic variation in components of dopamine neurotransmission impacts ventral striatal reactivity associated with impulsivity. *Molecular Psychiatry 14*, 60-70.

Kim, M. J., Scult, M. A., Knodt, A. R., Radtke, S. R., d'Arbeloff, T. C., Brigidi, B. D., & Hariri, A. R. (2018). A link between childhood adversity and trait anger reflects relative activity of the amygdala and dorsolateral prefrontal cortex. *Biological Psychiatry: Cognitive Neuroscience and Neuroimaging, 3*, 644-649.

Tan, H.-Y., Chen, Q., Goldberg, T. E., Mattay, V. S., Meyer-Lindenberg, A., Weinberger, D. R., & Callicott, J. H. (2007). Catechol-O-methyltransferase Val158Met modulation of prefrontal-parietal-striatal brain systems during arithmetic and temporal transformations in working memory. *Journal of Neuroscience, 27*, 13393-401.

Wager, T. D., & Nichols, T. E. (2003). Optimization of experimental design in fMRI: A general framework using a genetic algorithm. *Neuroimage, 18*, 293-309.

Zeineh, M. M., Engel, S. A., Thompson, P. M., & Bookheimer, S. Y. (2003). Dynamics of the hippocampus during encoding and retrieval of face-name pairs. *Science, 299*, 577-580.

**Supplementary Tables**

Supplementary Table 1. Brain regions whose intrinsic functional connectivity with the left basolateral amygdala showed significant positive association with trait anger ($p < 0.05$, corrected for multiple comparisons across the whole brain).

| Brain Region | Side | *Z* | *x* | *y* | *z* | # of voxels |
|---|---|---|---|---|---|---|
| *Seed: Left Basolateral Amygdala* | | | | | | |
| Putamen | R | 4.12 | 16 | 14 | -2 | 10362 |
| | L | 4.01 | 16 | 12 | -2 | |
| Caudate | R | 3.74 | 18 | 12 | 16 | |
| | L | 3.77 | -18 | 8 | 16 | |
| Nucleus Accumbens | R | 3.11 | 6 | 12 | -2 | |
| | L | 3.07 | -6 | 12 | -2 | |
| Pallidum | R | 3.83 | 18 | -2 | -6 | |
| | L | 3.55 | 18 | 0 | -2 | |
| Thalamus | R | 3.53 | 20 | -20 | 8 | |
| Amygdala | R | 3.01 | 24 | 0 | -22 | |
| Dorsomedial Prefrontal Cortex | R | 4.69 | 6 | 58 | 34 | 6729 |
| Dorsal Anterior Cingulate Cortex | R | 3.22 | 6 | 30 | 18 | |
| Inferior Frontal Gyrus | R | 3.50 | 52 | 14 | -2 | 236 |
| Inferior Temporal Gyrus | L | 4.47 | -64 | -18 | -30 | 87 |
| Middle Frontal Gyrus | L | 3.29 | -36 | 4 | 44 | 20 |
| Temporal Pole | R | 3.94 | 46 | 8 | -44 | 15 |
| Superior Frontal Gyrus | R | 3.73 | 6 | 12 | 70 | 12 |
| Orbitofrontal Gyrus | R | 3.54 | 40 | 22 | -18 | 11 |

*Note*: Coordinates are in Montreal Neurological Institute (MNI) space

Supplementary Table 2. Brain regions whose intrinsic functional connectivity with the right basolateral amygdala showed significant positive association with trait anger ($p < 0.05$, corrected for multiple comparisons across the whole brain).

| Brain Region | Side | *Z* | *x* | *y* | *z* | # of voxels |
|---|---|---|---|---|---|---|
| *Seed: Right Basolateral Amygdala* | | | | | | |
| Inferior Frontal Gyrus | L | 4.42 | -38 | 16 | 20 | 13140 |
| Dorsomedial Prefrontal Cortex | R | 4.12 | 8 | 48 | 40 | |
| Dorsal Anterior Cingulate Cortex | R | 2.90 | 2 | 26 | 24 | |
| Lateral Frontal Pole | R | 3.60 | 46 | 56 | 6 | |
| Medial Frontal Pole | R | 3.68 | 8 | 64 | -6 | |
| Supplementary Motor Area | R | 3.40 | 6 | -12 | 72 | |
| Lateral Occipital Cortex | R | 3.81 | 46 | -64 | 44 | 4036 |
| | L | 4.18 | -38 | -64 | 56 | 1335 |
| | L | 3.82 | -62 | -66 | 16 | 123 |
| Inferior Temporal Gyrus | R | 4.15 | 64 | -18 | -30 | 860 |
| Middle Temporal Gyrus | R | 3.39 | 68 | -48 | 0 | 501 |
| | L | 3.94 | -48 | -6 | -26 | 201 |
| Brainstem | L | 3.74 | -10 | -20 | -22 | 443 |
| | R | 4.74 | 10 | -20 | -22 | 128 |
| Cerebellum | R | 4.02 | 2 | -56 | -46 | 425 |
| Lateral Frontal Pole | L | 3.45 | -36 | 60 | -4 | 346 |
| | L | 2.56 | -46 | 46 | 6 | 22 |
| Caudate | L | 3.03 | -18 | 14 | 14 | 266 |
| | R | 3.36 | 14 | 14 | 0 | 204 |
| Superior Temporal Gyrus | L | 3.67 | -70 | -34 | 4 | 215 |
| Posterior Cingulate Cortex | R | 3.84 | 10 | -34 | 32 | 173 |

| | | | | | | |
|---|---|---|---|---|---|---|
| Thalamus | R | 3.22 | 8 | 0 | 10 | 162 |
| Supramarginal Gyrus | L | 3.05 | -68 | -22 | 22 | 130 |
| Hippocampus | R | 3.48 | 20 | -8 | -26 | 40 |
| Pallidum | L | 2.94 | -14 | -8 | -2 | 17 |
| Thalamus | L | 2.86 | -10 | 0 | -6 | 16 |
| Amygdala | R | 3.32 | 26 | 0 | -22 | 11 |

*Note*: Coordinates are in Montreal Neurological Institute (MNI) space

Supplementary Table 3. Brain regions whose intrinsic functional connectivity with the right centromedial amygdala showed significant positive association with trait anger ($p < 0.05$, corrected for multiple comparisons across the whole brain).

| Brain Region | Side | *Z* | *x* | *y* | *z* | # of voxels |
|---|---|---|---|---|---|---|
| *Seed: Right Centromedial Amygdala* | | | | | | |
| Inferior Frontal Gyrus | L | 5.57 | -36 | 16 | 20 | 22285 |
| Dorsomedial Prefrontal Cortex | R | 4.10 | 10 | 48 | 38 | |
| Dorsal Anterior Cingulate Cortex | R | 2.51 | 4 | 30 | 32 | |
| Lateral Frontal Pole | R | 3.36 | 50 | 56 | 4 | |
| Medial Frontal Pole | R | 3.56 | 8 | 66 | -6 | |
| Thalamus | R | 3.76 | 12 | -24 | 4 | |
| Caudate | R | 2.88 | 10 | 6 | 10 | |
| | L | 2.59 | -16 | 8 | 16 | |
| Amygdala | L | 3.37 | -20 | -6 | -12 | |
| Nucleus Accumbens | R | 2.96 | 12 | 16 | -4 | |
| Lateral Occipital Cortex | L | 3.67 | -38 | -72 | 50 | 342 |
| | L | 3.44 | -46 | -62 | 28 | 158 |
| | L | 4.02 | -62 | -66 | 16 | 63 |
| Orbitofrontal Cortex | L | 3.37 | -26 | 22 | -26 | 277 |
| Middle Temporal Gyrus | L | 3.64 | -50 | -42 | -2 | 263 |
| | R | 3.86 | 73 | -29 | 1 | 146 |
| | L | 3.12 | -58 | -52 | 0 | 15 |
| Temporal Pole | L | 3.35 | -58 | 10 | 0 | 144 |
| Superior Temporal Gyrus | R | 3.24 | 54 | -8 | -6 | 120 |
| Supplementary Motor Area | R | 3.53 | 14 | -6 | 42 | 119 |
| Inferior Temporal Gyrus | R | 4.49 | 66 | -18 | -28 | 94 |

| | | | | | | |
|---|---|---|---|---|---|---|
| | R | 3.50 | 60 | -54 | -20 | 14 |
| Inferior Frontal Gyrus | R | 3.33 | 60 | 18 | -2 | 44 |
| Precentral Gyrus | L | 3.47 | -54 | 0 | 20 | 37 |
| Posterior Cingulate Cortex | R | 3.35 | 10 | -34 | 32 | 37 |
| Heschl’s Gyrus | L | 2.84 | -42 | -24 | 2 | 25 |

*Note*: Coordinates are in Montreal Neurological Institute (MNI) space
